# The great acceleration of island saturation by species introductions in the Anthropocene has altered species-area relationships

**DOI:** 10.1101/2023.01.10.523426

**Authors:** Jason M. Gleditsch, Jocelyn E. Behm, Matthew R. Helmus

## Abstract

The species-area relationship (SAR) is a fundamental pattern of island biogeography which is often curvilinear due to reduced accumulation of species on mid-sized island caused by island isolation and the lack of speciation present on larger islands. The curvature of SARs represents lower saturation of species on mid-sized islands and therefore accelerated species accumulation should linearize island SARs. In the Anthropocene, island species accumulation has accelerated from introduced species. We hypothesize three new patterns. First, the saturation of species for the most unsaturated islands should increase more from introduced species than other islands. Second, SARs should become more linear as islands accumulate more species. Third, introduced species should greatly accelerate the island saturation process. We assessed these patterns for the reptile and amphibian of the Caribbean, a global hotspot of biodiversity. Mid-sized Caribbean islands are now more saturated causing a linearization of contemporary herpetofauna SARs resulting from a ca. 30 myr and 40 myr acceleration of island saturation for reptiles and amphibians, respectively. Thus, humans within the last few hundreds of years—starting with European colonization of the Americas—have greatly accelerated the natural process of island saturation by 30 million years within the Caribbean global biodiversity hotspot.

## Introduction

Islands naturally accumulate species over long, geologic timescales through over-water immigration and *in situ* speciation. Both are theorized to be related to geographic area and isolation^1–3^. Theory based on the principals of classic island biogeographic theory and subsequent theoretical advancements^1,2,4^ suggests that species accumulation increases the species richness of islands until available habitat (or niche space) is filled with species, a point referred to as the saturation point^5^. Given the positive relationship between island area and habitat diversity, and area’s direct effect on accumulation, saturation points are largely thought to be set by island area^1,6,7^ with factors limiting species accumulation, such as isolation reducing immigration, causing islands to be further from saturation. Isolated islands naturally have low immigration rates^1,3^, and thus, species accumulation rate is reduced and more reliant on in *situ* speciation. However, in *situ* speciation is positively influenced by island area, and thus islands without speciation have reduced accumulation. Therefore, an island’s level of saturation is dependent on how area and isolation influence immigration and speciation.

On small islands, the saturation point is low. There is little habitat diversity on small islands and even without *in situ* speciation, immigration is often sufficient to saturate habitats if the islands are not too isolated. As island area increases, the saturation point increases, but speciation (specifically, cladogenic *in situ* speciation) does not occur until the islands are large enough for species to differentiate^8^. The lack of speciation on smaller islands causes islands to be increasingly further away from their saturation point as area increases with mid-sized islands without speciation the furthest from saturation^8^. This creates weak species-area relationships (SARs) for small to mid-sized islands^8^. When islands are large enough to support speciation, the accumulation of species is increased and the relationship between species richness and area is stronger^8–10^. Theory suggests that the reduced saturation of mid-sized islands without speciation causes log-log SARs across islands of a biogeographic region to be curvilinear^11–14^. If species accumulation greatly accelerates, SARs should become more linear over time with mid-sized, isolated islands quickly approaching their area-defined saturation points (Fig. 1).

**Figure 1:**
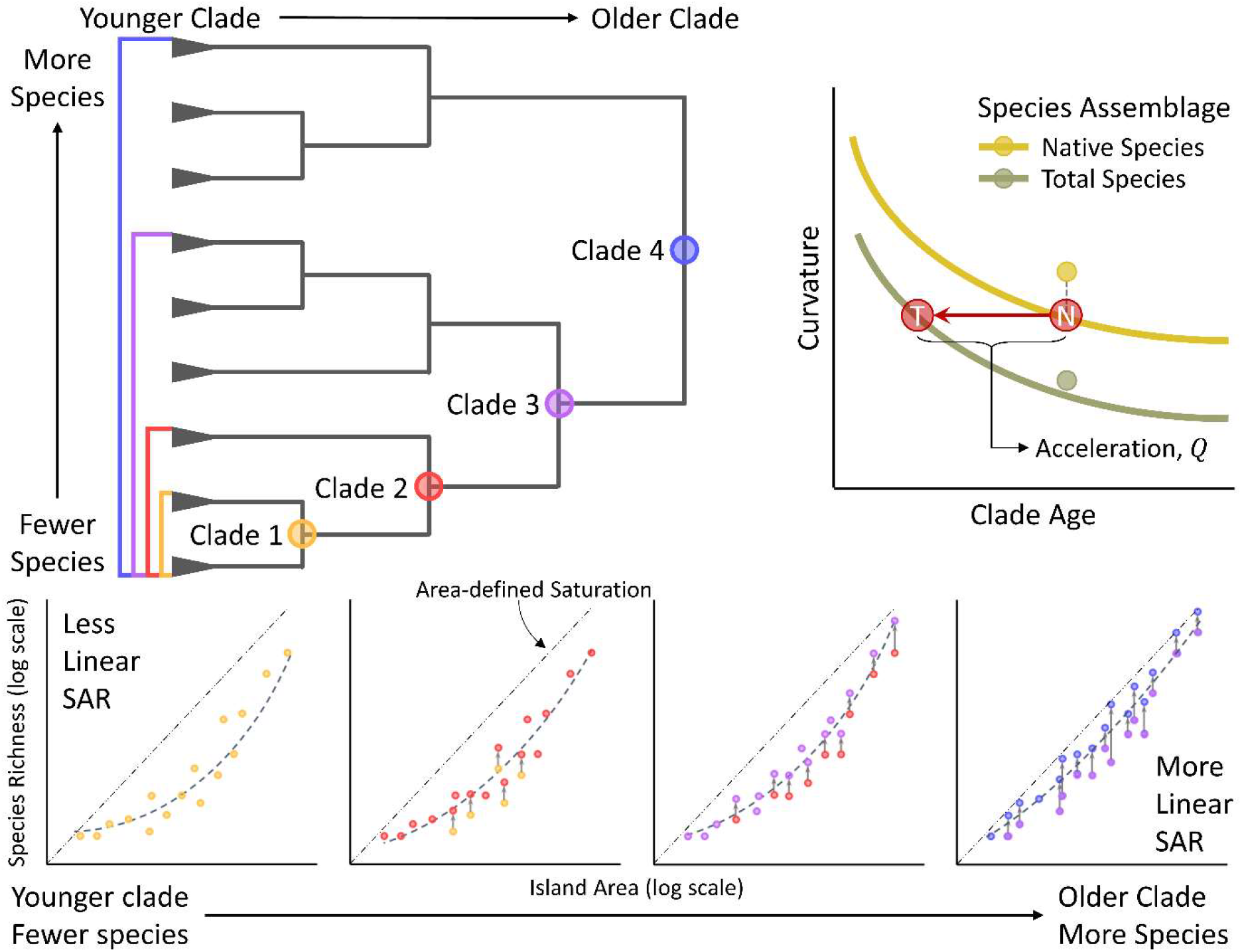
Phylogenetic scale is expected to influence the curvature of species-area relationships (SARs). As scale increases, SAR curvature should decrease because more time has passed for species to accumulate on islands. (top left) A phylogenetic tree with four clades of species (colored circles and brackets) delineated with Clade 1 at the smallest phylogenetic scale and Clade 4 at the largest phylogenetic scale. (bottom) SARs naturally become more linear as phylogenetic scale increases. Gray arrows represent changes in species richness from the previous phylogenetic scale (points colored as the previous scale). The increased saturation of islands as phylogenetic scale increases is depicted by how the islands (points) approach their saturation points (area-defined saturation line). Because of increasing saturation, the species-area relationships become more linear as phylogenetic scale increases. The greatest increase is for the mid-sized islands that linearizes the SAR as scale increases (see main text for reason). (top right) The increase in saturation due to introduced species decreases the time it takes to linearize SARs allowing for the estimation of island saturation acceleration by comparing the relationship SAR curvature has with clade age between native (gray) and total (yellow) species assemblages.

In the Anthropocene, species accumulation rates are accelerating worldwide due to the immigration of introduced species through human activity^15^ that connects islands with distant regions and new species pools^16^. Given the increased accumulation rate and species pool size in the Anthropocene, we hypothesize, based on the principals of classic island biogeographic theory and subsequent theoretical advancements^1,2,4^, the emergence of three new biogeographical patterns resulting from species accumulation and saturation. First, mid-sized islands that are furthest from being saturated with species should have a greater change in their level of saturation due to introduced species than the islands with closer to saturation. Second, as unsaturated mid-sized islands become more saturated, contemporary SARs that include both native and introduced species should be more linear than SARs that include just native species (a proxy for past SARs). Third, since introduced species have greatly accelerated island immigration, the island saturation process should also be greatly accelerated in the Anthropocene. We term these three hypothesized phenomena: increased island saturation, species-area relationship linearization, and acceleration of island saturation.

Here, we used reptile and amphibian (i.e., herp) species in the greater Caribbean region to test for evidence of our three hypothesized biogeographic patterns. To address the *increased island saturation* hypothesis, we first described the relationship island saturation has with isolation, area, and speciation to support the theory that the least saturated islands are the mid-sized and isolated islands. Then, we tested if banks least saturated with native species had greater changes in their level of saturation due to introduced species than the islands with greater native species saturation. To address the *species-area relationship linearization* hypothesis, we estimated how SAR curvature differs among three assemblages of species based on whether they are native or introduced across all herp clades. These three assemblages were comprised of native, introduced, or total (native + introduced) species. Specifically, if the SARs for the total species assemblages were more linear than SARs for the native assemblages (proxy for past SARs) and the most linear SARs were for the introduced species assemblage, then contemporary SARs (i.e., total species assemblage) are more linear due to the increased immigration from species introductions. To address the *acceleration of island saturation* hypothesis, we tested how the saturation of Caribbean islands changed over time for the native and total species assemblage by leveraging the temporal scales of the herp phylogenetic tree (i.e., phylogenetic scale; see top left in Fig. 1 and Methods). For this, we expect SARs to become more linear as phylogenetic scale increases. It is expected from theory that at larger phylogenetic scales islands have more time to become saturated (bottom row in Fig. 1; see Methods). Therefore, we determined the relationship between SAR curvature and phylogenetic scale (proxy for time) along lineages of nested clades for native and total species assemblages. Then, we quantified the acceleration of island saturation in the Anthropocene by estimating the difference between the SAR curvature-time relationships of native species and total species assemblages (top right in Fig. 1).

## Results and Discussion

The accumulation rate of herp species in the Caribbean region has accelerated by six orders of magnitude compared to the past accumulation of native species (Extended Data Fig. 1). Our records of Caribbean herp species included 1039 extant native (N = 1006) and introduced (N = 95; 32 species from outside the Caribbean) amphibian and reptile (i.e., herps) species from 171 genera on 72 island banks (i.e., island groups based on historical land connections and underwater topography) that ranged in size from 0.119km^2^ to over 110,000km^2^. Here, we will refer to island banks as islands for simplicity.

### Has accumulation rate greatly accelerated for Caribbean herps?

We estimated the difference in accumulation rates of native and introduced species by comparing the past accumulation rate of native species endemic to the Caribbean islands to the contemporary accumulation rate of introduced species. We found that past accumulation was a million times slower than contemporary accumulation. For past accumulation, based on a lineage through time plot^17^ of Caribbean endemics (see Methods), a new species was added naturally to the Caribbean approximately every one million years (1.016 × 10^−6^ species/yr; Extended Data Fig. 1A). In contrast, the accumulation rate of species introduced into the Caribbean was approximately one species added every year (1.011 species/yr; Extended Data Fig. 1B). This acceleration in accumulation rate was also observed for individual islands with accumulation rates between five and eight orders of magnitude higher past accumulation (Extended Data Table 1). There are two caveats to this comparison. First, lineage through time plots do not incorporate extinct lineages, but given the difference in rates, the number of extinct lineages would have astronomically high to reasonably affect our results. Second, it is unknown if all currently established introduced species will persist in the long term. Regardless, the accumulation rate of species of the Caribbean has greatly accelerated in the Anthropocene, even though our estimate lacks exact precision.

### Increased Island Saturation

The main factors influencing the natural level of saturation for Caribbean herps were island isolation and the presence of *in situ* speciation. The least saturated islands were the most isolated and mid-sized islands that lacked *in situ* speciation (Fig. 2). This was determined by developing a metric of saturation by taking the native and total species assemblage residuals from a 95% quantile regression of the natural log of total species richness on the natural log of island area (see Fig. 2A). Like the observed negative species-isolation relationship (Fig. 2B) that is predicted by island biogeography theory ^1^, we found that more isolated islands were less saturated with native species than less isolated islands when the effect of area and the presence of speciation were controlled (Fig. 2C). Mid-sized islands were less saturated in native species as expected with curvilinear SARs (Extended Data Table 2). This drop in native saturation on mid-sized islands was explained by island isolation (Fig. 2C), island area, and the presence of speciation on the island (Fig. 2D; Extended Data Table S) as expected by theory ^8,10^. Therefore, the curved SAR observed for native Caribbean herps (see Fig. 2A) was the result of the combined effects of island isolation, which limits immigration, and island area, which limits speciation. This is further evidenced by the steep relationship between area and saturation when speciation is not present on banks and the lack of relationship when speciation is present (Fig. 2D). When islands have speciation, their species richness is increased by both immigration and speciation giving rise to higher species accumulation rates (Extended Data Table 1).

**Figure 2:**
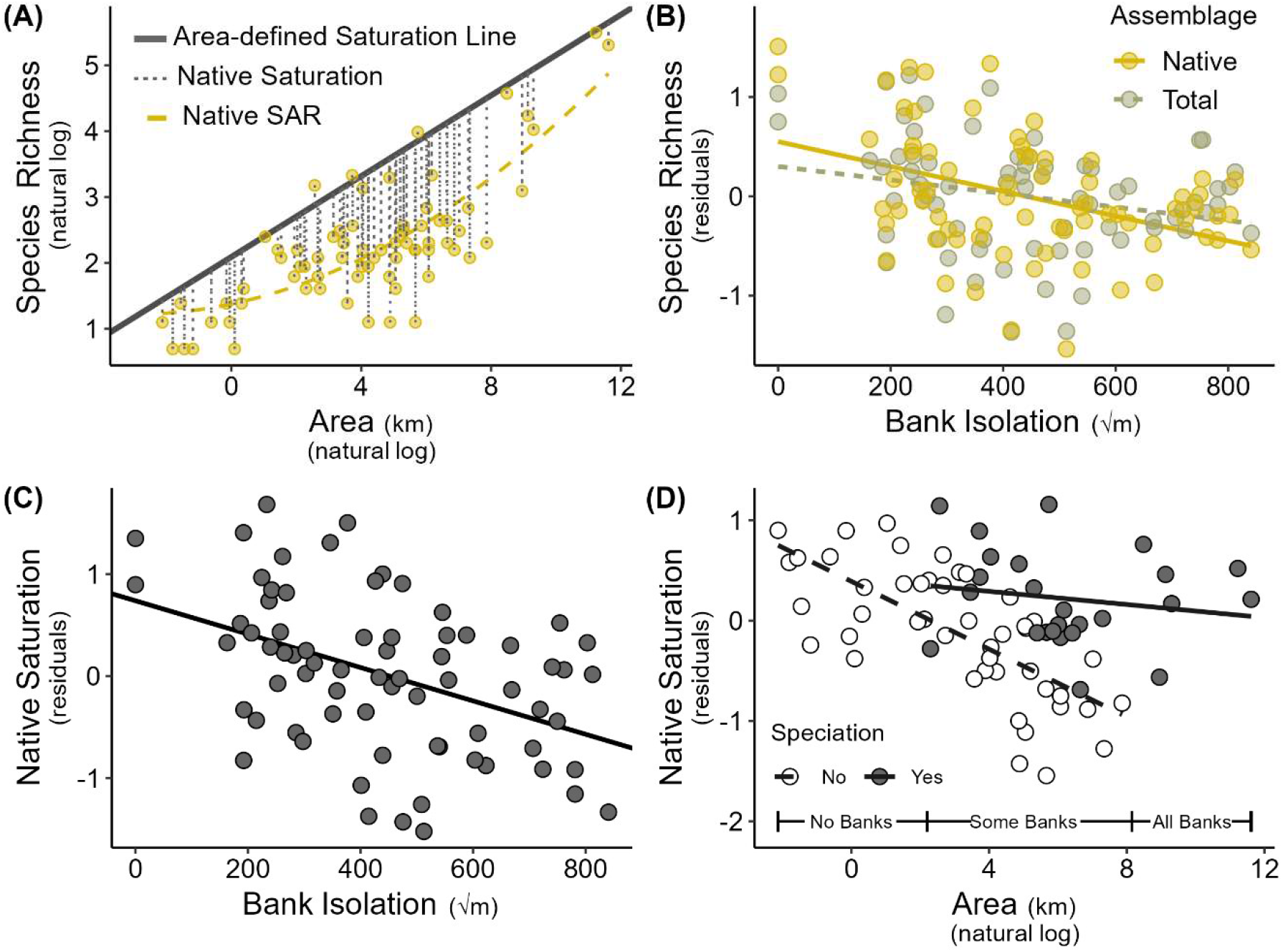
For all the Caribbean reptiles and amphibians, the island banks that were the most isolated and had no speciation were the least saturated with native species. As expected by island biogeography theory, these effects were most pronounced in mid-sized banks. (A) The saturation of banks with native species was estimated as the difference between the natural log of their species richness and their area-defined saturation point estimated by a 95% quantile regression for the total (native + introduced) species assemblage (dark gray line). (B) More isolated banks had lower species richness. To control for the effect of area the residuals from the relationship between species richness and area are plotted. (C) The more isolated banks had lower saturation of native species when the effects of area and speciation on native saturation were controlled (plotted are the residuals from the relationship between native saturation and area, speciation, and their interaction). (D) The saturation of banks with speciation (filled points, solid trendline) had no relationship with bank area, but saturation decreased with area for banks without speciation (unfilled points, dashed trendline). To account for the effect of isolation (see C), the residuals from the relationship between native saturation and isolation were used in D. Along the x-axis in D is a scale representing the amount of speciation across the banks in three categories of bank area based on the number of banks in the area range that have speciation (see Methods).

As a result of species introductions, Caribbean islands are becoming more saturated. The highest change in saturation on the islands that are less saturated with native species, as expected (Fig. 3). We calculated the change in saturation due to introduced species as the difference between native saturation and total saturation (Fig. 3A). The change in saturation due to introduced species decreased as islands were more saturated with native species (Standardized Estimate = −0.519, SE = 0.102, *t*-value = −5.077, *p*-value < 0.001; Fig. 3B), suggesting that the least saturated islands approached their area-defined saturation point more than the islands with higher saturation. Further, as expected, the islands that exhibited the most change in their saturation also exhibited a stronger negative relationship between their change in saturation and native saturation (75% and 25% quantile regression estimates: −0.840 and −0.305, respectively). However, since isolation influences species accumulation and therefore decreases native saturation^1,3^, the more isolated islands should also have a greater increase in their level of saturation due to introduced species than less isolated islands. Indeed, the more isolated islands were more saturated from the introduction of species than less isolated islands (Standardized Estimate = 0.428, SE = 0.111, *t*-value = 3.844, *p*-value < 0.001; Fig. 3C).

**Figure 3:**
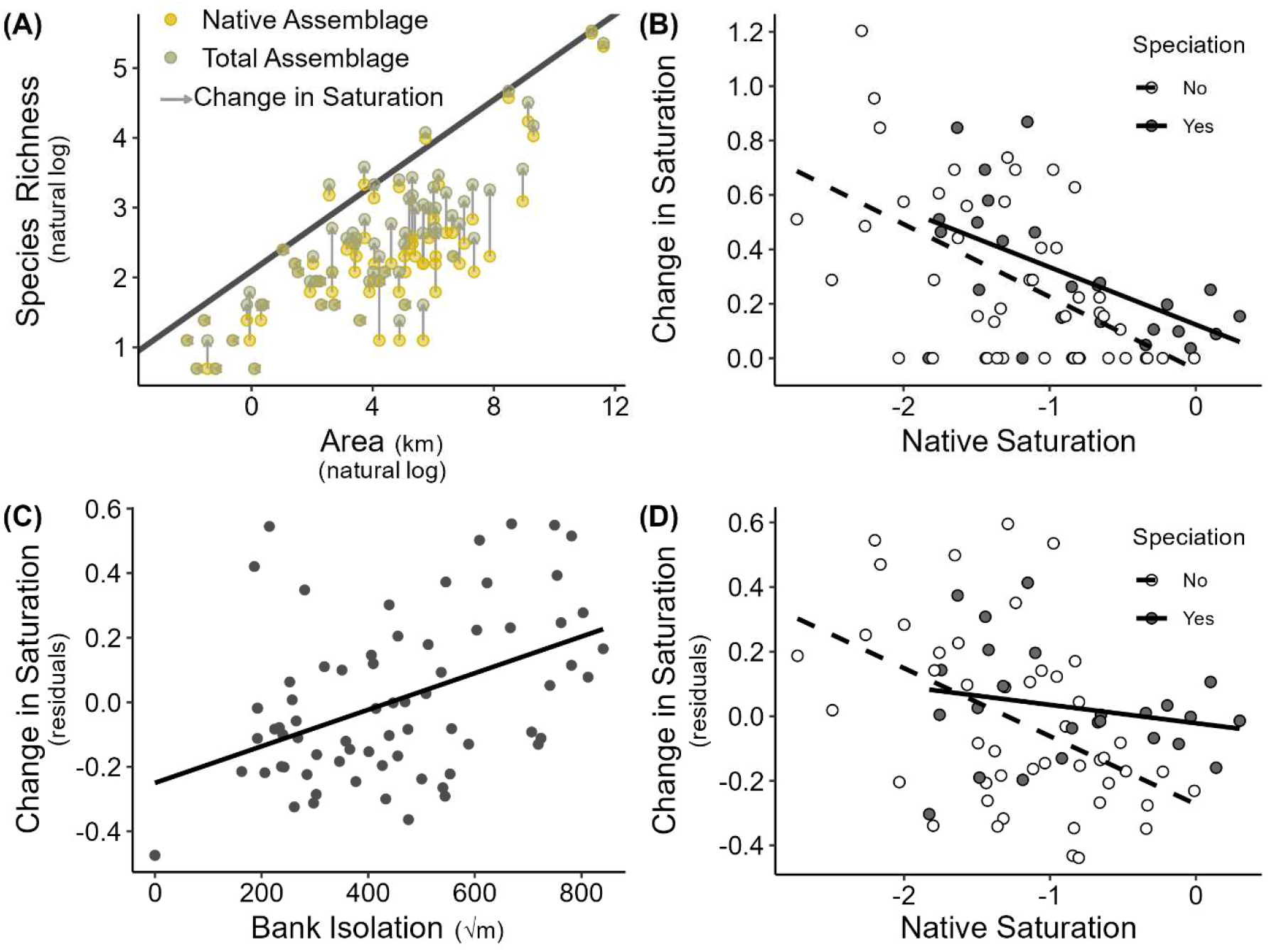
In the Anthropocene, banks least saturated with reptile and amphibian species increased in saturation more than banks naturally closer to saturation. This change in saturation was influenced by both isolation and speciation. (A) Island banks had their species richness (native species richness = yellow points) increased by the anthropogenic introduction of species (total species richness = gray points) causing them to approach their estimated area-defined saturation point (dark gray line). (B) The change in saturation due to introduced species was greater the less saturated a bank was for both banks without speciation (unfilled points, dashed trendline) and banks with speciation (filled points, solid trendline). (C) The change in saturation due to introduced species was also greater the more isolated a bank was when the effect of area was accounted for (plotted are the residuals from the change in saturation–bank area regression). (D) When the effect of isolation was accounted for (residuals from the change in saturation–bank isolation regression), the effect of native saturation on the change in saturation due to species introductions was greatly reduced on banks with speciation suggesting the main driver of the negative relationship in C for banks with speciation is bank isolation.

Our results are congruent with other studies that have shown that the more isolated and species-poor islands gain more species than less isolated, species-rich islands from the introduction of species^14,18^. When the effect of isolation is controlled for by regressing the residuals of the change in saturation–isolation relationship on native saturation, we see that the negative relationship observed between the change in saturation and native saturation for islands with speciation (filled points and solid line in Fig. 3B) is driven largely by island isolation (Fig. 3D). The increased accumulation of species due to introductions also weakened the expected species-isolation relationship (Fig. 2B) likely due to a breakdown of traditional geographic barriers to immigration^14^.

### Species-Area Relationship Linearization

As expected, Caribbean herp SARs are now more linear than in the past (Fig. 4). Across 59 phylogenetic clades, a curved model fit native SARs better than a linear model suggesting that Caribbean herp SARs are curved based on the adjusted R^2^ of the curvilinear vs linear models (Fig. 4A). Although, the increased ability of a curved model to approximate the species-area relationship for the total species assemblage was less than for the native species assemblage (Fig. 4A). To further investigate the shape of the SARs, we developed a curvature metric, 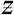, by calculating the area between the first- and second-order regression lines of the SARs (see Methods). This metric allowed us to statistically test the degree to which the introduction of species has linearized the SARs of Caribbean herps. The SARs of both the introduced and total species assemblages were significantly more linear, in terms of the curvature metric, than the native species assemblage (mean percent differences = 73.54 and 28.37, respectively; Fig. 4B; Extended Data Table 3); a result that was consistent across other metrics of curvature (see Methods). The increased linearity of the SARs for the total species assemblages when compared to native assemblages suggests a linearization of the herp SARs in the Caribbean due to anthropogenic species introductions.

**Figure 4:**
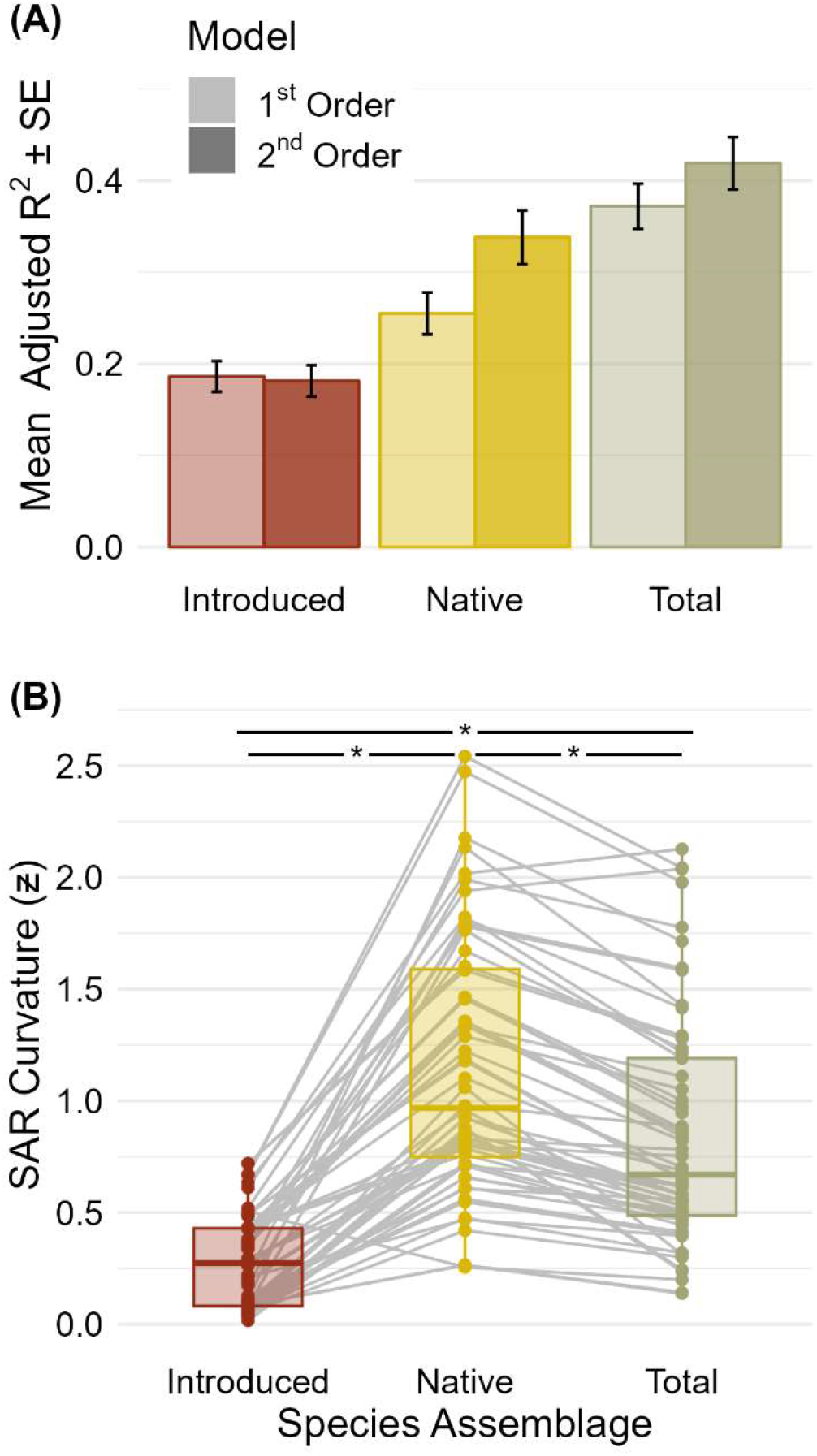
The contemporary species-area relationships (SARs) of amphibian and reptile clades in the Caribbean have been linearized due to species introductions. (A) The second order (i.e., curved) SARs generally fit better for native and total species assemblages across all clades but that difference was higher for the native than the total species assemblage. Shown are the average (± s.e.) adjusted R^2^ for the first and second order (lighter and darker color, respectively) SARs across all clades. The higher difference in fit for the native species assemblage suggests more curved SARs, which was supported by the curvature metric. (B) Boxplots of the SAR curvatures 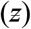 of the three species assemblages (gray lines connect SAR curvatures of the same clade to show the pairwise comparisons) show the differences in curvature between the species assemblages. All differences in curvature between the assemblages were significant at α = 0.01 according to a paired Wilcoxen sign-rank test as shown by the stars and lines at top of plot (see Extended Data Table 3 for statistics).

The hypothesized linearization of SARs due to increased immigration from introduced species was further supported by the observed patterns of SAR curvature across the different species assemblages on banks (Fig. 4B). Across all clades, the assemblage that had the most linear SARs was that of only introduced species. The linear SARs of introduced species suggests that isolation and traditional geographic barriers to dispersal have little influence on the spread of introduced species (as suggested by ^15,18^). This conclusion is further supported by our observation of larger increases in species saturation on isolated, species-poor islands (Fig. 3)^18^.

Thus, the more linear SARs of the total species assemblages (i.e. the assemblages that contain introduced and native species) is likely from a reduction in the influence of isolation and increased immigration from the human-mediated introduction of species^14,19^. Theory suggests that isolation should reduce the rate of immigration to banks^1^ causing the slow process of *in situ* speciation to be the main driver of species accumulation on isolated islands. However, since isolation is no longer a strong influence on immigration rate, mid-sized islands can accumulate more species faster bringing them closer to their area-defined saturation point.

### Acceleration of Island Saturation

Given enough time, theory suggests islands will accumulate species and approach their area-defined species saturation points^2,20^ creating a more linear SAR across an island system^14^. To determine the relationship between time (i.e., clade age) and SAR curvature, we focused on the reptile and amphibian evolutionary lineages. Starting with the two largest herp genera in the Caribbean, *Anolis* lizards and *Eleutherodactylus* frogs, and working backwards along the lineages to Reptilia and Amphibia, we calculated SAR curvature for all nodes, delineating clades, along the lineages. Both evolutionary lineages exhibited SARs that became more linear as the clades became older and larger (Fig. 5A and B). To better understand the isolated effects of clade age and size we performed a bootstrapping analysis (see Methods) and found that clade age had a stronger effect than clade size suggesting the primary driver of the SAR linearization with phylogenetic scale is the age of the clade (Fig. 5 insets).

**Figure 5:**
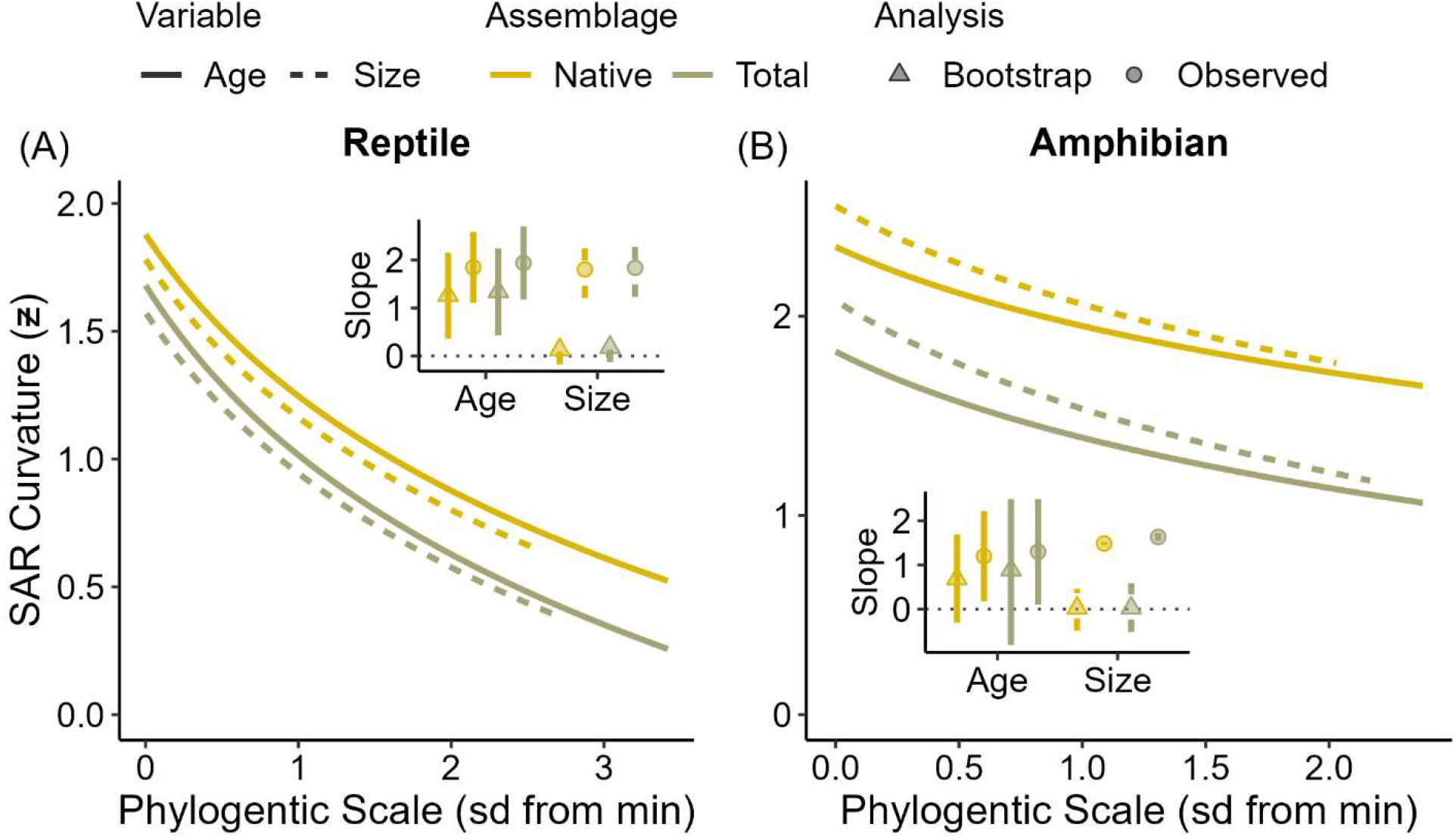
Amphibian and reptile clades in the Caribbean at larger phylogenetic scales exhibited more linear species-area relationships (SARs). The curvature of the SARs along both the (A) Reptile (*Anolis*) and (B) Amphibian (*Eleutherodactylus*) lineages decreased as phylogenetic scale (standard deviations from minimum) increased irrespective of whether scale was measured by clade age (solid line) or clade size (dashed line). This was also true for both the native (yellow) and the total (gray) species assemblages. The fixed-age bootstrapping (triangles with solid line in inset) of the relationship between curvature and clade age for both the Reptile and Amphibian lineages also exhibited negative relationships. However, the fixed-clade size bootstrapping (triangles with dashed line in inset) of the relationship between curvature and clade size did not exhibit a strong negative relationship and was weaker than the bootstrap relationship with clade age. This suggests the observed negative relationship with phylogenetic scale is due primarily to the increasing of clade age with phylogenetic scale, not increased species pool size. The x-axis in both plots is the number of standard deviations from the minimum of clade age and size to represent phylogenetic scale.

However, based on theory, the accumulation of species should slow with time as islands become increasingly saturated (i.e., approach maximum species richness) and species fill available habitat^2,5^ so the linearization of SARs should slow with time. This pattern was previously found with diversification rates of Caribbean *Anolis* lizards on the four Greater Antillean islands such that rates slowed down with time suggesting islands become saturated with species given enough time^21^. The linearization of SARs we observed along evolutionary lineages followed a logarithmic curve with the rate of linearization decreasing as SAR curvature approached zero (Fig. 5). This supports the idea that species accumulation rates slow as islands approach saturation and that investigating the shape of SARs across phylogenetic scale can be viewed as analogous to the island saturation process. Our analyses support this idea and extend it to include the accumulation of species not only through diversification but also immigration and introduction.

Through their effect on immigration rate, we estimate that humans have accelerated the island saturation process by millions of years (Fig. 6). Since the difference between the relationships of clade age and SAR curvature for native and total species assemblages is due to the introduction of species by humans, we were able to observe and estimate this rescaling by comparing these relationships. Our results showed that island saturation has been sped up by between 12.35 and 65.43 myr for reptiles (*Anolis* lineage) and between 33.14 and 127.97 myr for amphibians (*Eleutherodactylus* lineage) with the median rescaling factor 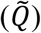 being approximately 33.70 and 43.07 myr, respectively (Fig. 6A and B). The acceleration of island saturation by tens of millions of years is a new example of how human activity is rescaling ecological and environmental processes in the Anthropocene.

**Figure 6:**
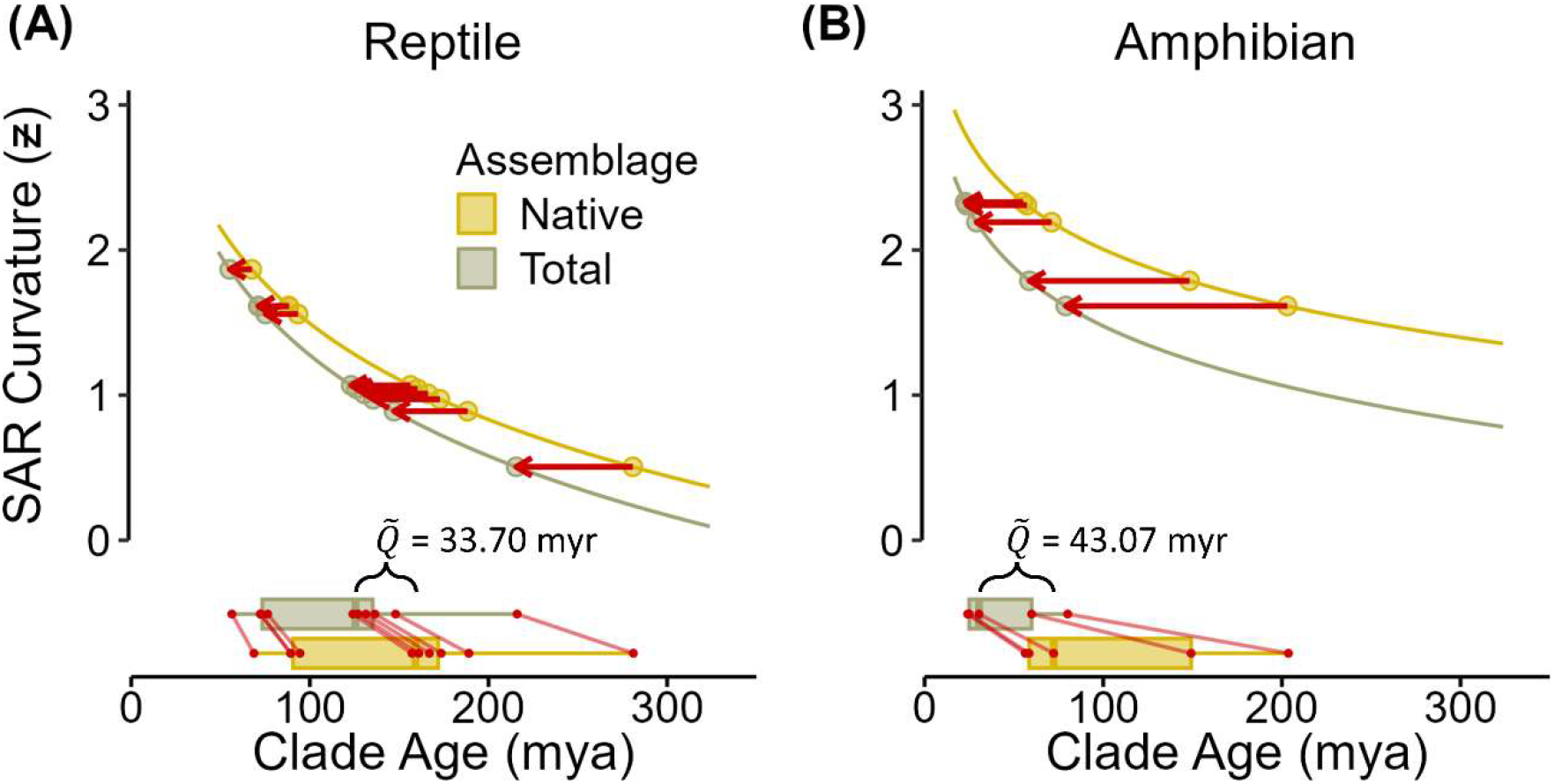
Humans have accelerated the island saturation process by millions of years. The difference between the relationship between phylogenetic scale and SAR curvature of native and total species assemblages suggests a rescaling of island saturation with species as determined by the difference in the predicted age of the total and native assemblages (red arrows and boxplots with median difference, 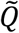) and this effect was less for the (A) Reptile (*Anolis*) than for (B) Amphibian (*Eleutherodactylus*) lineage. The fitted value of 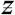 of the native assemblage for each clade was projected onto the 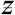 – clade age curve of the total species assemblage, and the difference between the two ages was used as the rescaling factor, *Q*. This was done for each clade and the median taken 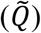 for an overall rescaling factor as shown by the box plots along the bottom (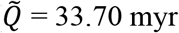 for reptiles and 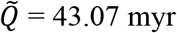 for amphibians).

As the world progresses through the Anthropocene, naturally occurring processes and patterns will become increasingly influenced by human activity giving rise to modified spatiotemporal scales at which these processes and patterns operate^22^. One such example of this is the temporal rescaling observed here of island saturation leading to the linearization of the species-area relationship. Currently, introductions are outpacing extinctions on islands suggesting that, globally, islands have not reach their area-defined saturation point^23,24^, and human-mediated introductions do not show signs of slowing down^14,25,26^ (Extended Data Fig. 2). The effect of human-mediated introductions on the shape of the SARs increased over time (Fig. 5 and Extended Data Fig. 4 and Table 5). That, along with increased effect of time on SAR curvature as accumulation and species pool size increases (Extended Data Fig. 5 and Table 4), suggests that human activity will have increasing effects on SAR curvature and island saturation as source pools expand and rates of introduction increases through economic trade.

If the rates of introduction and extinction shift so that extinctions become more frequent, the speed at which islands are becoming saturated in the Anthropocene will likely result in native species making up a disproportionate number of extinctions. This is because native species will have little time to adapt to the increasing pressures of novel competitors and predators as well as having less habitat to sustain large populations due to the conversion of natural land cover for anthropogenic purposes. While the land development can initially increase habitat diversity by opening new habitat into which introduced species can immigrate^27^, as development continues, habitat diversity is likely to decrease leading to reductions in species richness^19^. Therefore, the relationship between area and habitat diversity may become decoupled as human activity increases on islands, which can further alter the area-defined species richness equilibrium of islands and influence the shape and strength of SARs. Together the increased immigration due to introduced species along with the extinction of native species would lead to an overall decrease in beta and gamma diversity of island systems across the globe. This should be of concern for biodiversity conservation since islands hold an exceptionally high amount of the world’s species, especially those that are threatened or endangered, given their land area^28^.

## Methods

### Study System

The greater Caribbean region is a global biodiversity hotspot that has high levels of endemism^29^ and is composed of mostly oceanic islands that exhibit increasing economic connections, making it an ideal region to investigate the effect of human activity on island biogeography^30^. The island groups included in the greater Caribbean region are the Greater and Lesser Antilles, Bahamas, Turks and Caicos, Southern Antilles, Trinidad and Tobago, Cayman Islands, and the Mexican, Belizean, Honduran, Columbian, and Nicaraguan Islands as defined by Hedges *et al*^31^. In addition, we included Bermuda as in Helmus *et al*.^14^ due to its high level of trade with the Caribbean and 75% of its reptile and amphibian species also occur in the Caribbean^19^ (see also ^32^). These islands were then grouped into island banks (referred to as islands in the main text) based on Powell and Henderson^33^ and the underwater topography (see Supplementary Information Fig. S1) obtained using the General Bathymetric Chart of the Oceans^34^. Island banks are clear macroevolutionary units that are statistically robust due to their separation by areas of deep ocean causing natural overwater dispersal among banks to be much less than dispersal within banks (e.g., ^14,31,35,36^).

### Bank Area and Isolation

To estimate the area-defined saturation line and determine the species-area relationships for the Caribbean herps, we calculated the area for each island bank by summing the land area of each island in the island bank. Island area was determined from the GADM shapefile (version 3.6^37^), projected into the Lambert azimuthal equal area projection (centered at 72° W, 20° N), using “st_area” function in the ‘sf’ R package^38^. The area of each island bank was then natural log transformed.

To determine if isolated island banks were less saturated with native species, we calculated the isolation of each bank from potential evolutionary source pools. Based on phylogeographic reconstructions of ancient dispersal pathways^39–41^ we considered mainland South and Central America plus Cuba and Hispaniola as natural source pools for Caribbean herps. The distance from the shorelines of each of these sources to the geographic centroid of each island bank was determined using the world GADM shapefile, projected into the azimuthal equidistant projection (centered at 72° W, 20° N), using “st_distance” function in the ‘sf’ R package. The square root of the minimum distance to an evolutionary source was used as our metric of isolation^42^.

### Species Occurrence Data

We limited our taxonomic scope to the vertebrate classes Reptilia and Amphibia (i.e. herps), which comprise 46% of the terrestrial vertebrate species globally^43^ and comprise at least 40% of the terrestrial vertebrate species in the Caribbean including approximately 75% of the endemic terrestrial vertebrate species^29^. The two most specious genera of Caribbean terrestrial vertebrates are *Anolis* lizards and *Eleutherodactylus* frogs^44^. We used a dataset of island-level herp records compiled from an extensive literature search to determine the native, introduced, and total (native + introduced) species richness for each island bank^19^ that was updated with new information.

The literature search was completed in June 2022. We primarily used Powell and Henderson^33^ to obtain lists of the species for most of the islands in the greater Caribbean region but needed to add records of species from islands not included in this publication. For these other islands, we obtained species records from field guides and books as well as and extensive literature search (see ^19^). For introduced species we used Hailey, Wilson and Horrocks^45,46^ and added to that introduced species records found through literature searches on Google Scholar, Web of Science, and natural history and occurrence notes of species checklists and Caribbean, herp, and invasion ecology journals. We also added records downloaded from Global Biodiversity Information Facility (GBIF)^47^ using the R package ‘rgbif’^48^ that were of “research grade” and of species that had multiple records spanning multiple years. The literature search and GBIF records yielded 1039 extant species, 95 of which have been introduced outside their native range, and 21 extinct species of herps on 72 banks.

Species taxonomy was standardized and their status (i.e., whether they were native or introduced on particular banks) were checked with caribherp.org^49^, Reptile Database^50^, Amphibian Species of the World 6.1^51^, the Integrated Taxonomic Information System (ITIS^52^, accessed June 2022), and GBIF.

### Species Accumulation in the Caribbean

We determined if species accumulation rate has increased during the Anthropocene by comparing the past native accumulation rate to the accumulation rate of introduced species from 1550 through 2018. To estimate past native accumulation rates, a lineage through time (LTT) analysis^17^ was used to calculate the slope of the lineage through time accumulation plot of the phylogeny of endemic Caribbean clades (see Extended Data Fig. 1A). The phylogeny used in the LTT analysis was obtained from a dated tetrapod phylogeny downloaded from timetree.org^53^ (accessed December 2022) that included 711 species – including 4 possibly extinct species – out of 1,053 total herp species. To determine the accumulation rate of introduced species, we calculated the slope of an accumulation curve (see Extended Data Fig. 1B) of species introduced from outside the Greater Caribbean region between 1150 and 2018 based on the earliest year a species was recorded in the literature or estimated to have been introduced. This comparison was done for the entire Caribbean region and repeated for each bank that had enough native species to calculate the slope of the LTT plot to see how bank-specific species accumulation has changed in the Anthropocene (see Extended Data Table 1). N PX FE F

### Species-area Relationship Curvature

To understand how the shape of species-area relationships (SARs) has changed in the Anthropocene, we created a metric to describe the curvature of SARs in way that allowed us to investigate the shape of SARs across clades and phylogenetic scales. Non-linear SARs often exhibit weak slopes for smaller to midsized islands, and strong slopes for midsized to large islands and are often represented by two-part regressions^11,13,14^ Simple first order regressions can describe the overall linear trends of SARs, but two-part SARs are better approximated with a second order regression. Therefore, the extent that the SARs are curved can be measured by the difference between the first order SAR regression (*f*(*A*); eq. 1) and the second order SAR regression (*g*(*A*); eq. 2), which we term SAR curvature, represented by 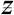. SAR cuvature is calculated by the definite integral of the absolute value of the difference between *f*(*A*) and *g*(*A*), where *A* is island area (eq. 3). With this metric, as 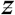 approaches zero, SARs are more linear.

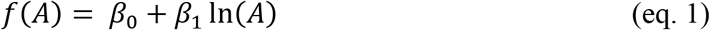

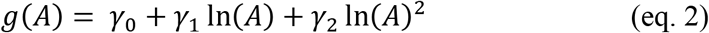

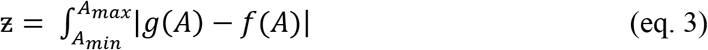

The increase in saturation due to introduced species can then be determined by the difference between the curvature of native and total SARs and is represented by percent reduction in 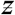 (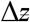, eq. 4).

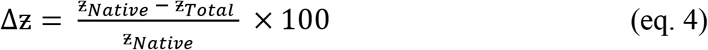

We validated 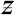 by testing its congruence to two alternative curvature metrics. The first alternative metric was based on model goodness of fit and was the absolute value of the difference between the residual sum of squares of *f*(*A*) and *g*(*A*). The second alternative metric of curvature was the parameter of curvature for *g*(*A*), which is calculated using equation 5, where *g*(*A*)’ and *g*(*A*)” are the first and second derivatives, respectively, of *g*(*A*). This parameter of curvature is then averaged across all values of bank area. Both these metrics show linearization as they approach zero, and our metric, 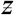, was highly congruent with both alternative metrics (Supplementary Information Fig. S2).

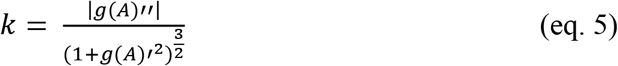

### Hypothesis 1 – Increased Island Saturation

To test our hypothesis that the saturation of isolated, species-poor island banks increased more than species rich island banks due to species introductions, we estimated how far the native and total species richness of each island bank was from the estimated area-defined saturation line (see Fig. 2A). Additionally, to see if island bank isolation had the predicted negative relationship with island bank species richness (species-isolation relationship, SIR), we regressed the residuals from the first-order species-area relationship (note: this is not the saturation metric but is similar) on bank isolation (Fig. 2B). We estimated the area-define saturation line by performing a 95% quantile regression between island bank area and total species richness (see ^14^) using the ‘rq’ function in the ‘quantreg’ R package^54^. Both area and total species richness were natural log transformed. The native and total saturation of each island bank was then estimated by calculating the difference between the area-defined saturation line and the natural log transformed native and total species richness (see Fig. 2A). To understand the effect isolation has on native saturation while controlling for the effects of area and speciation, we regressed the residuals from the relationship native saturation has with bank area, the presence of speciation, and their interaction on bank isolation (Fig. 2C). Then, to understand the effect of area on native saturation while controlling for the effect of isolation, we regressed the residuals from the relationship native saturation has with isolation on bank area. The relationship between area and isolation was broken up between island banks with speciation and island banks without speciation (Fig. 2D). The change in saturation was calculated as the difference between the estimated total (i.e., contemporary) and native (i.e., past) saturation of each island bank (see Fig. 3A). The change in saturation was then regressed on native saturation and island bank isolation to determine if isolated, species-poor island banks had the most change in their saturation (see Fig. 3A and B). In order to understand the mechanism behind this relationship between native saturation and the change in saturation due to introduced species we regressed the residuals of the change in saturation–island bank area relationship on island bank isolation (Fig. 3C). Then we used the residuals of the change in saturation–island bank isolation relationship and regressed those on native saturation to see how the relationship changes when the effect of isolation is controlled (Fig. 3D). To explore how the effects of native saturation and island bank isolation are different depending on the amount of change observed, we also performed a 75% and 25% quantile regression.

### Hypothesis 2 – Linearization due to Species Introductions

To test whether SARs are more linear due to species introductions, we compared the curvature of SARs, 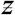, between the native, introduced, and total (native + introduced) species assemblages of 57 herp clades. We formed the clades by grouping species based on their phylogenetic relationships determined from the dated tetrapod phylogeny. First, we removed all non-herp species and treated genus as the smallest clade (e.g., Clade 1 in Fig. 1). Then tracing each branch that originated from the node of each genus’s most recent common ancestor, we grouped species into sequentially older and larger clades at every node until we came to the root of the tree with two clades, one for all reptiles and one for all amphibians (see Fig. 1). This process resulted in 71 reptile clades and 22 amphibian clades. However, some of these clades were found only on a limited number of island banks. Thus, we only used distinct clades and retained only those clades that were found on enough banks to have high statistical power to detect area relationships based on a power analysis using the fit of an all-herp second order SAR (R^2^ = 0.70; numerator DF = 2; power = 0.8). The final number of clades used in our analysis were 43 reptile and 16 amphibian clades.

We regressed the log transformed species richness (i.e. ln(SR + 1)) on the log transformed bank area (i.e. ln(Area)) for the first order SAR (*f*(*A*), eq. 1) and added a square term for area for the second order SAR (*g*(*A*), eq. 2) for the native, introduced, and total assemblages of each clade. This produced six separate SARs (i.e., three first order and three second order) for each clade. We standardized (mean centered and divided by the standard deviation) both the transformed species richness and island bank area to make the curvature metric comparable between clades.

Then, we compared the curvature of SARs, 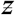, between each pair of the species assemblages for each clade using a paired Wilcoxon sign rank test. We calculated 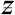 for each assemblage of each clade and tested the hypothesis that SARs for total species assemblages are more linear than the SARs for native species assemblages. Additionally, we tested if the introduced species assemblage had the most linear SARs compared to the native and total assemblages since they would be the most linear if isolation had little effect of isolation on the human-mediated immigration of introduced species.

Given that these clades are not completely independent, we repeated these analyses with just the terminal clades (n = 11) and used phylogenetic linear mixed models (PLMM) to compare the assemblages while accounting for the phylogenetic relationships of the clades. We performed the PLMMs using the ‘pglmm’ function in the “phyr” R package^55^. The results of the PLMMs were congruent with our results from the Wilcoxon sign rank tests (Extended Data Table 3) and had *p*-values less than 0.0001 (Extended Data Table 6).

### Hypothesis 3 – Acceleration of Island Saturation

To test our hypothesis that island saturation has accelerated due to species introductions, we determined how island saturation has changed through time for native and total herp species assemblages in the Caribbean. Testing how island saturations changes through time is difficult given the long, geologic timescales on which species accumulation operates. For species introduced to islands through human activity, species accumulation can be directly estimated, but this is not possible for native species. However, for native species that arose on islands through colonization or speciation, a dated phylogeny can be leveraged to estimate time when investigating biogeographical patterns. An example of this is the use of Dynamical Assembly of Islands by Speciation, Immigration and Extinction (DAISIE) models to estimate biogeographic rates of archipelagos and has been used to determine equilibrium dynamics and how they are influenced by human-induced extinctions (e.g., ^3,56,57^). Here we employ the use of phylogenetic scale to investigate the saturation of islands across timescales.

Phylogenetic scale is the position of clades relative to each other in a phylogeny (top left in Fig. 1) and can be characterized by the age of the common ancestor of all species in the clade (i.e., crown age in millions of years before present) and the size of the clade (i.e., number of species that descended from the common ancestor)^58^. Both clade age and size increase with phylogenetic scale as one moves from the present to the past along an evolutionary lineage in a dated phylogeny (i.e., Clades 1 through 4 in Fig. 1). Thus, phylogenetic scale is related to time such that as phylogenetic scale increases so does the time for species accumulation via colonization and *in situ* speciation. Given that at larger phylogenetic scales there is more time for species accumulation, we would expect for islands in a region to be more saturated with species, and therefore exhibit more linear SARs, at larger phylogenetic scales (Fig. 1).

First, we determined how SAR curvature of the of clades at varying phylogenetic scales to test the theorized relationship between phylogenetic scale and SAR curvature and thus the use of phylogenetic scale as a proxy for the island saturation process. This was done along the evolutionary lineage of the two largest genera in the Caribbean, *Anolis* lizards (reptile) and *Eleutherodactylus* frogs (amphibian). These lineages allowed us to use nested clades needed for this type of analysis (see Fig. 1) while keeping enough statistical power at lower phylogenetic scales to estimate robust SARs. For this, we regressed 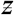 on the natural logarithm of clade age (i.e., crown age in millions of years before present) for each lineage and then also on the natural logarithm clade size (i.e., number of species that descended from the common ancestor). Again, all variables in these models were standardized. Since the clades at smaller phylogenetic scales are nested within clades at larger scales along evolutionary lineages, we compared generalized least squares regressions with a first order autoregressive (AR1) covariance structure to regressions without an AR1 structure using Akaike information criterion for small sample sizes (AICc). This comparison showed that an autoregressive structure is not needed for our analyses. For the visualization of these relationships, we transformed clade age and clade size into z-scores centered on the minimum value (i.e., the number of standard deviations from the minimum) to represent phylogenetic scale irrespective of how it was measured (see Fig. 5).

Since both clade age and size increase with phylogenetic scale, we performed a fixed-age and -size bootstrapping analyses to isolate the effects of clade age and size. By doing this we were able to determine how much the effect of phylogenetic scale on SAR curvature was due to clade age versus clade size, and therefore, support our conclusion that decreasing curvature with phylogenetic scale represents species saturation over time. First, to isolate the effect of clade age, we randomly sampled 50 species from each clade along the reptile (*Anolis*) and amphibian (*Eleutherodactylus*) lineages 1,000 times (referred to as fixed-size bootstrapping). We selected the 50 species so that the age of the clade was the same as it was observed (e.g., the 50 species of the Reptilia clade always had a crown age of ca. 281myr). We then regressed the SAR curvature of these random clades against the natural logarithm of the clade age. To isolate the effect of clade size, we then randomly sampled groups of species of varying size (240 – 500 species by 10 for the *Anolis* and 160 – 220 species by 10 for the *Eleutherodactylus* lineages) from the clade that had a crown age of ca. 173.5myr for the *Anolis* lineage and ca. 72.3myr for the *Eleutherodactylus* lineage 1,000 times keeping the clade age constant (referred as fixed-age bootstrapping). We then regressed the SAR curvature of these random clades against the natural logarithm of the clade size. In both lineages the effect of age when clade size was held constant was stronger than the effect of clade size when age was held constant (Fig. 5) suggesting that the effect of phylogenetic scale on SAR curvature was more due to increasing clade age than clade size.

Therefore, the effect of species introductions on island saturation can be determined by the difference of the relationships clade age has with SAR curvature for the native and total species assemblages. To see if this effect is greater at larger phylogenetic scales, we regressed 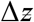 on clade age and then on clade size. If 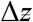 increases with both clade age and size, then it is likely the acceleration of island saturation is due to the compounding effects of increasing species pool size and higher species accumulation rates. To further support the idea that increase species pool sizes and accumulation rates drive the difference between native and total assemblages, we separated the reptile clades into groups of species based on if they arose on banks through *in situ* speciation, natural immigration, or human-mediated introduction. To do this we performed an ancestral state reconstruction for the bank of origin for species in the herp phylogeny. The traits that were put into the reconstruction were the island banks of endemic species or mainland for species with mainland populations, and the likelihood of each island bank, or mainland, was determined for each ancestral node. The ancestral state reconstruction was done using an equal-rates model with the ‘ace’ function in the ‘ape’ R package^59^. For each species that was not endemic to an island bank, the most likely island bank or mainland of the ancestral node was used. We then calculated SAR curvature for the species the arose through speciation, then for both speciation and immigration, and then for all species (i.e., now including introduced species) and regressed these curvature values on clade age. Since the species accumulation rate and the available species pool increases with each group, if the effect of clade age on SAR curvature increases (i.e., increased magnitude of slope) with each step, the proposed mechanism of compounded effects of species pool size and increased accumulation is further supported.

Finally, to estimate the amount that species introductions have accelerated island saturation in the Anthropocene, we calculated the difference between the fitted values of 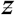 for native assemblage at an observed clade age and the clade age for the equal fitted value of 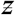 for the total species assemblages, *Q* (top right in Fig. 1). The was done for each clade along the *Anolis* and *Eleutherodactylus* lineages and the median of those values was used for the rescaling estimate, 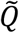 (Fig. 6).

## Acknowledgements

We would like to thank the members of the Integrative Ecology Lab at Temple University for their comments during the conception and early draft of this work. JMG, JEB, and MRH were supported by Temple University.

