## Extended Data for "The great acceleration of island saturation by species introductions in the Anthropocene has altered species-area relationships"

### Figures

**Figure 1**: The accumulation rate of introduced species in the Caribbean is much higher than native species. Shown in (A) is the lineage accumulation through time plot of reptile and amphibian species endemic to the Caribbean with the slope (in years instead of myr) of lineage accumulation displayed (log scale; red dashed line). Shown in (B) is the accumulation of introduced reptile and amphibian species through time with the slope of the curve displayed (log scale; red dashed line). Note: The x-axis in (A) is millions of years while the x-axis in (B) is in years.

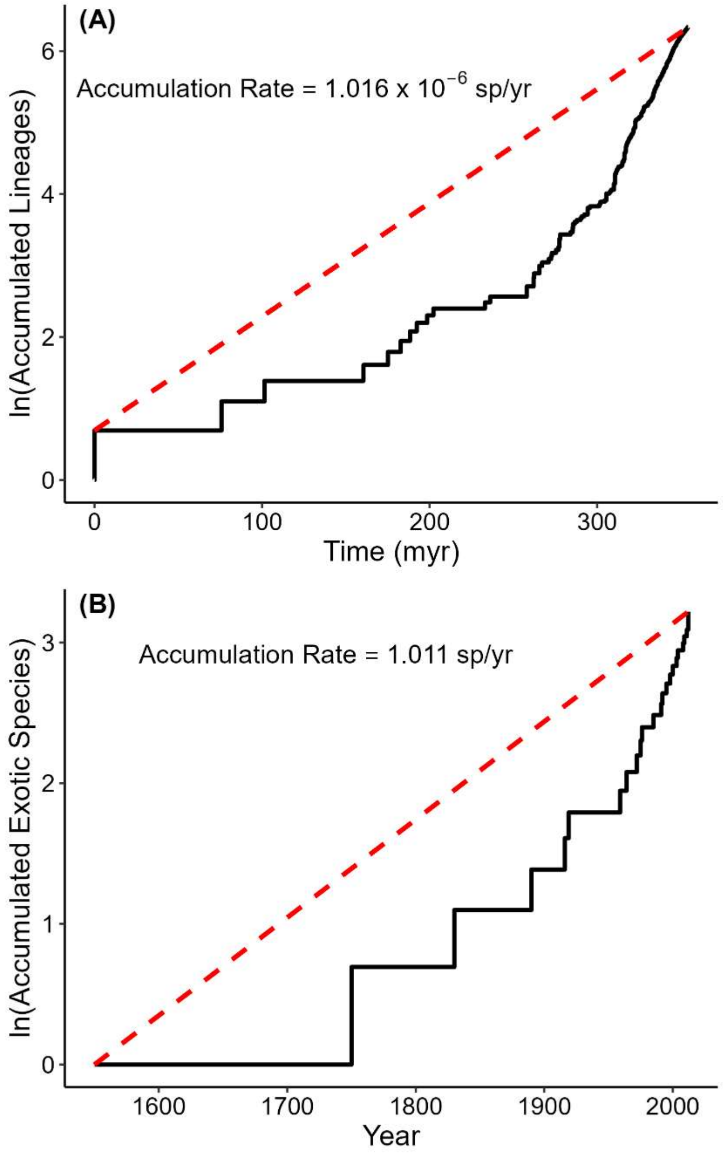

**Figure 2**: The acceleration of species introductions in the Caribbean. Shown in the accumulative number of introduction events (black points fitted with a LOESS black line; left y-axis) and the decadal introduction rate (red points fitted with a LOESS red line; right y-axis) as determined by dividing the number of introduction events that occurred in a decade by ten years. Figure adapted from Gleditsch et al. (2022)[19].

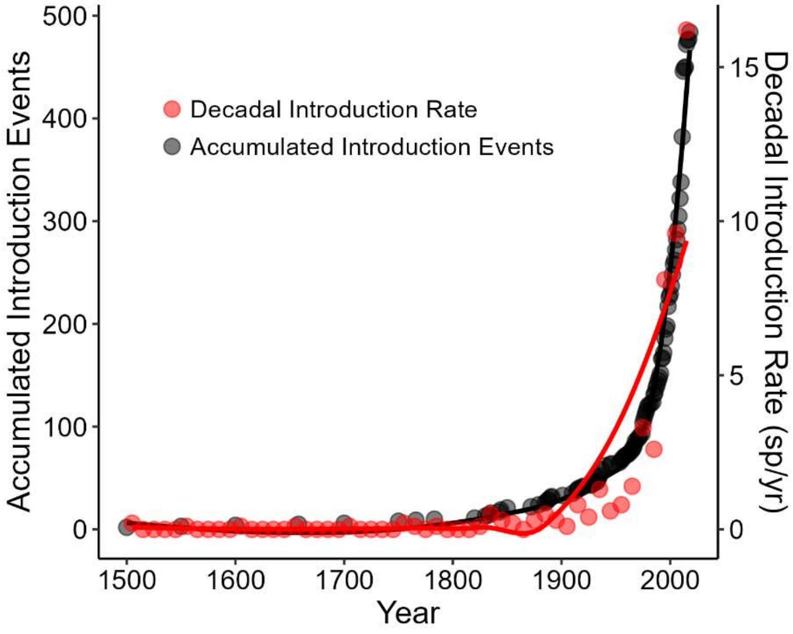


**Figure 3**: The linearization effect of introduced species increased with both clade age (solid line, circles) and clade size (dashed line, triangles) along the (A) Reptile (*Anolis*) and the (B) Amphibian (*Eleutherodactylus*) lineages. The x-axis in both plots is the number of standard deviations from the minimum of clade age and size to represent phylogenetic scale.

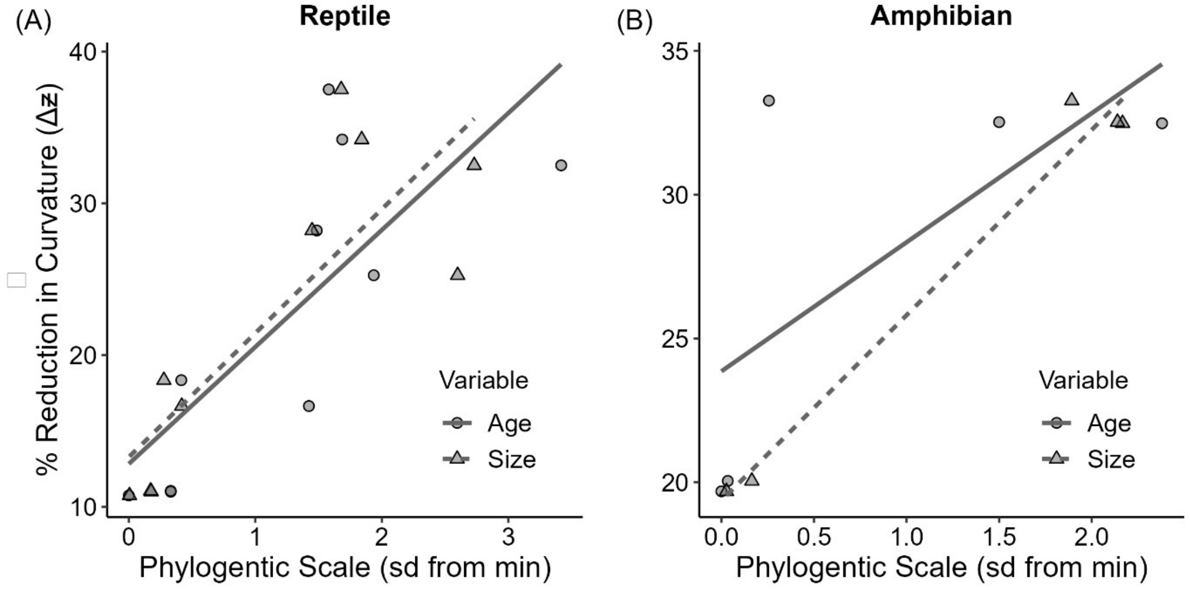

**Figure 4**: The effect of phylogenetic scale measured by clade age increased as the rate of species accumulation increased from just speciation (tan) to speciation and colonization (orange) to speciation, colonization, and introduction (dark brown). These groups also exhibit an increase in geographic region from which species accumulate: from just the island (speciation) to the island and the rest of Caribbean bioregion (speciation and colonization) to the island, Caribbean bioregion, and global trade partners (speciation, colonization, and introduction). Species origins on islands were determined by distribution and an ancestral state reconstruction (see Methods).

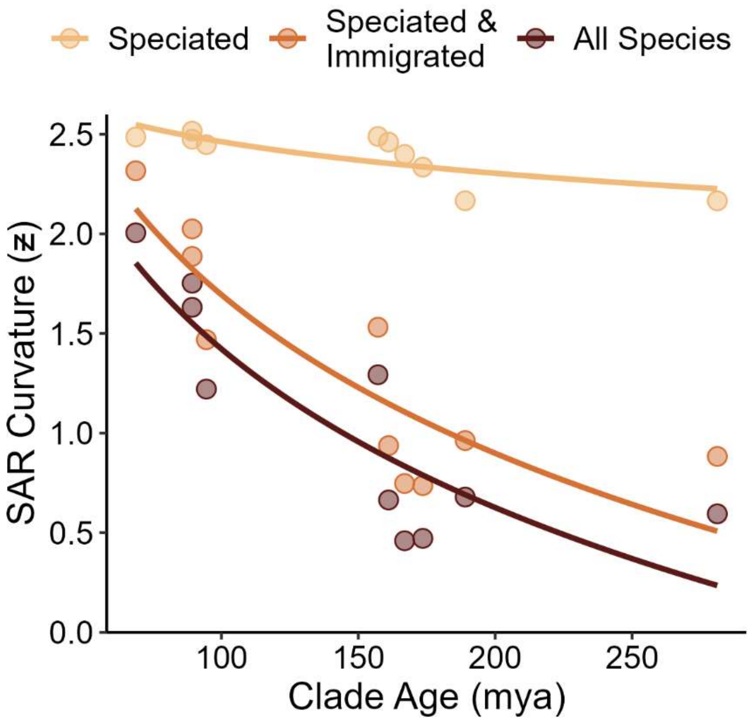

**Tables**

**Table 1**: Each bank wth enough species for rate estimation had introduction rates 5 to 9 orders of magnitude greater than past species accumulation rates. Shown are the rates of the introduced and endemic species accumulation as estimated by species accumulation and lineage through time plots. The rates have been transformed to represent the increase in ln(Species) per year. Stars represent the banks that were estimated to have speciation. The bolded rows are banks with the highest estimated speciation rates that also had the highest accumulation rates of endemics aside from Trinidad and Tobago which are large banks right off the South American coast.

| Bank | Accumulation Rate of Introductions | Endemic Accumulation Rate | Orders of Magnitude Difference |
|---|---|---|---|
| **Anguilla*** | **0.0096** | **$5.70 \times 10^{-09}$** | **6** |
| Antigua* | 0.0143 | $3.10 \times 10^{-09}$ | 7 |
| **Cuba*** | **0.0076** | **$1.20 \times 10^{-08}$** | **5** |
| Dominica* | 0.0180 | $3.10 \times 10^{-09}$ | 7 |
| Grenada* | 0.0269 | $3.10 \times 10^{-09}$ | 7 |
| Guadeloupe* | 0.0092 | $3.50 \times 10^{-09}$ | 6 |
| **Hispaniola*** | **0.0096** | **$1.22 \times 10^{-08}$** | **5** |
| **Jamaica*** | **0.0050** | **$8.30 \times 10^{-09}$** | **6** |
| Martinique | 0.0055 | $2.60 \times 10^{-09}$ | 6 |
| Montserrat | 0.0825 | $4.00 \times 10^{-09}$ | 7 |
| **Puerto Rico*** | **0.0355** | **$9.20 \times 10^{-09}$** | **7** |
| St. Lucia* | 0.0157 | $4.70 \times 10^{-09}$ | 7 |
| St. Vincent* | 0.0029 | $2.00 \times 10^{-09}$ | 6 |
| Tobago* | 0.3277 | $2.00 \times 10^{-09}$ | 8 |
| Trinidad* | 0.0075 | $8.70 \times 10^{-09}$ | 6 |

**Table 2**: Area had a curved relationship with the native saturation of banks with lower saturation on midsized banks. The curved relationship was explained by bank isolation and the presence of speciation on the bank as evidenced by the non-significant square term of area when isolation and speciation were added to the model.

| Explanatory Variable | Estimate | Standard Error | *t*-value | *p*-value |
|---|---|---|---|---|
| Area | -0.275 | 0.109 | -2.520 | 0.014 |
| Area^2 | 0.228 | 0.079 | 2.882 | 0.005 |
| Area | -0.800 | 0.157 | -5.090 | < 0.001 |
| Area^2 | -0.041 | 0.107 | -0.381 | 0.704 |
| Speciation | 0.867 | 0.229 | 3.777 | < 0.001 |
| Isolation | -0.378 | 0.086 | -4.401 | < 0.001 |
| Area:Speciation | 0.734 | 0.363 | 2.023 | 0.047 |

**Table 3**: Comparisons of SAR curvature from native, introduced, and total species assemblages show the highest curvature is in the native assemblage and the most linear SARs are the introduced species assemblages. Shown are the mean differences and the test statistic and *p*-value from a paired Wilcoxen rank tests for each pairwise comparison.

| Comparison | Mean Difference | Test Statistic | *p*-value |
|---|---|---|---|
| Native-Total | 0.304 | 17 | < 0.0001 |
| Native-Introduced | 0.875 | 3 | < 0.0001 |
| Total- Introduced | 0.571 | 18 | < 0.0001 |

**Table 4**: Results from linear models of $z$ (natural log transformed) regressed separately on clade age and clade size for each species assemblage of the *Anolis* and *Eluethrodactylus* lineages. Also shown are the estimates for the age-curvature relationship when the Anolis lineage is broken up to those species that arose on islands through speciation, speciation and immigration, and all species (includes introduced species).

| Explanatory Variable | Lineage | Assemblage | Estimate | Standar d Error | *t*-value | *p*-value |
|---|---|---|---|---|---|---|
| Clade Age | *Anolis* | Native | - 0.261 | 0.060 | - 4.381 | 0.002 |
| | | Total | - 0.346 | 0.080 | - 4.311 | 0.003 |
| | *Eluethrodactylus* | Native | - 0.138 | 0.058 | - 2.368 | 0.099 |
| | | Total | - 0.203 | 0.093 | - 2.186 | 0.117 |
| Clade Size | *Anolis* | Native | - 0.722 | 0.121 | - 5.962 | < 0.001 |
| | | Total | - 0.933 | 0.159 | - 5.852 | < 0.001 |
| | *Eluethrodactylus* | Native | - 1.294 | 0.006 | - 207.580 | < 0.001 |
| | | Total | - 1.884 | 0.060 | - 31.295 | < 0.001 |
| Clade Age | *Anolis* | Speciated | -0.074 | 0.022 | -3.359 | 0.010 |
| | | Speciated & Immigrated | -0.629 | 0.137 | -4.584 | 0.002 |
| | | All Species | -0.813 | 0.185 | -4.382 | 0.002 |

**Table 5**: Results from linear models of the percent change in curvature ($z$) between native and total species assemblages regressed separately on clade age and clade size (total species richness) for each species assemblage of the *Anolis* and *Eluethrodactylus* lineages. All relationships were significant at the α = 0.05 level.

| Lineage | Explanatory Variable | Estimate | Standard Error | *t*-value | *p*-value |
|---|---|---|---|---|---|
| *Anolis* | Clade Age | 0.853 | 0.185 | 4.262 | 0.002 |
| | Clade Size | 0.903 | 0.152 | 5.951 | < 0.001 |
| *Eluethrodactylus* | Clade Age | 0.744 | 0.386 | 1.926 | 0.150 |
| | Clade Size | 0.993 | 0.070 | 14.117 | < 0.001 |

**Table 6**: Results of a phylogenetic generalized mixed model testing if the SARs of the 11 terminal clades were more linear for the introduced species assemblage than the total and native assemblages, and whether the SARs of the total assemblage were more linear than the native assemblage. The larger the estimate the more curved the SAR of the second assemblage in the comparison. The results that matched the Wilcoxen rank tests of all the clades are represented by a star in the "Matched" column.

| Assemblage Comparison | Estimate | Std. Error | *z*-score | *p*-value | Matched |
|---|---|---|---|---|---|
| Introduced-Total | 0.8299 | 0.1926 | 4.3080 | < 0.0001 | * |
| Introduced-Native | 1.1561 | 0.2227 | 5.1920 | < 0.0001 | * |
| Total-Native | 0.3262 | 0.0748 | 4.3596 | < 0.0001 | * |
