## Supplementary Information for "The great acceleration of island saturation by species introductions in the Anthropocene has altered species-area relationships"

**Figure S1**: Map of the Greater Caribbean region with underwater topography shown in shades of blue and the banks outlined in red. Map taken from Gleditsch et al. (2022).

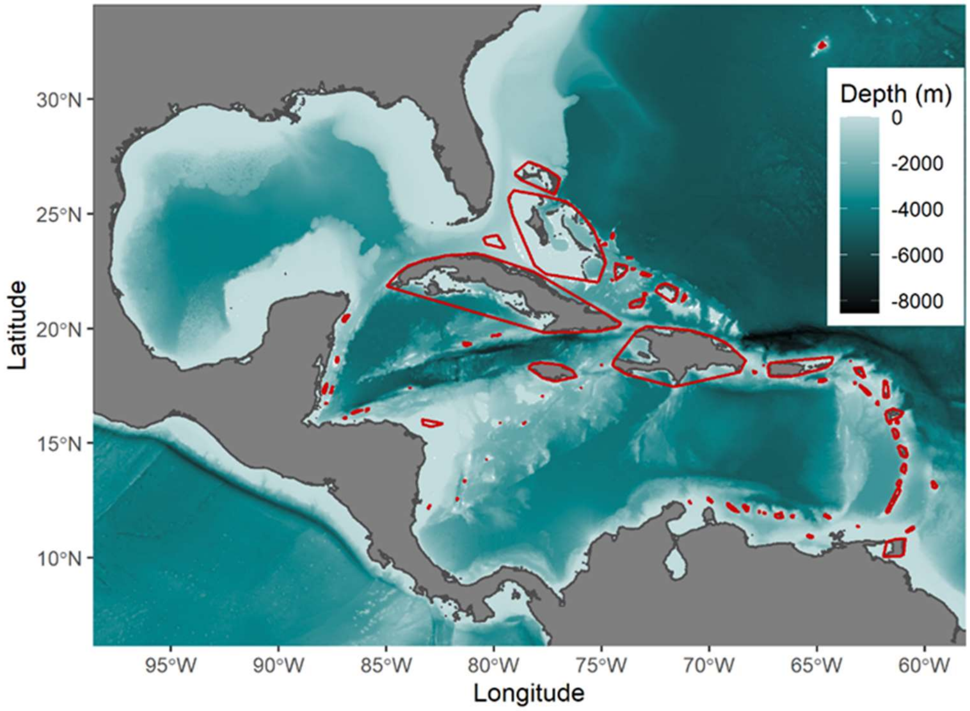


**Figure S2**: The correlation between the three measures of SAR curvature were all above 0.9 suggesting that $z$ is a good metric of SAR curvature.

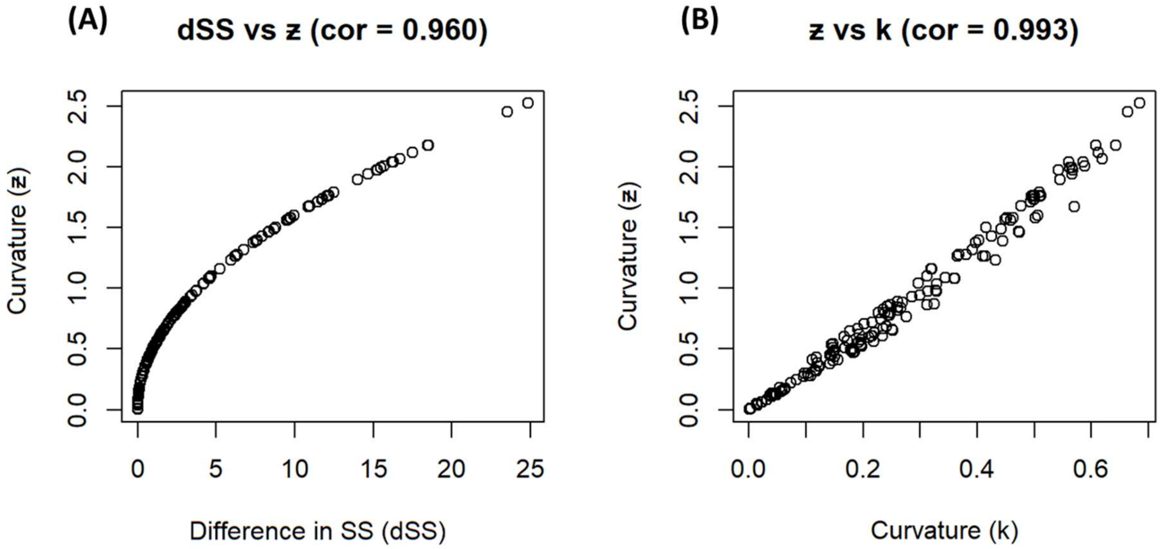
